# A demographic model for estimating the inter-division lifespans of stem cells and the subsequent transit amplifying stages

**DOI:** 10.1101/521336

**Authors:** Purna Gadre, Shambhabi Chatterjee, Bhavna Varshney, Debasmita Mazumdar, Samir Gupta, Nitin Nitsure, Krishanu Ray

## Abstract

The division rates of stem cells and their progeny shape the growth and maintenance of tissues. Here, we present a mathematical model that could estimate the stage-wise lifespans of germline stem cells (GSCs) and subsequent transit amplifying (TA) cells from their steady-state distribution in *Drosophila* testis. Analysis of the wild-type data using this model indicated that the inter-division lifespans of the first two TA cycles remain similar to that of the GSCs, and then reduce by nearly 2-folds for the third and fourth cycles. Also, loss of Cyclin E and Cdk1 functions in the early germline cells, which decreased the rates of GSC divisions, is suggested to extend the lifespans of GSCs and the TA stages without affecting subsequent differentiation. Similar perturbations at the 4 and 8-cell stages, however, arrested the mitoses at the 8-cell stage, and only the Cyclin E-deficient cells continued with premature meiosis. Together, these results suggest that regulation of the G1-S and G2-M transitions in the GSCs and the rapidly dividing TA stages differentially impacts the amplification of the germline pool and subsequent differentiation. The model also helped to quantify distinct influences of these cell cycle regulatory molecules in determining the lifespans at different TA stages.

**Highlights:** A model for calculating the lifespans of transit amplifying stages from demography. Transit-amplifying divisions accelerate by nearly 2-folds after the second mitosis. Cyclin E and Cdk1 regulate the lifespans of GSCs and transit amplifying cells. The premature arrest of the final transit amplifying division induces meiosis.

## Introduction

Regulation of stem cell divisions is critical during development for tissue growth and during adulthood for tissue maintenance. All stem cells have the ability to undergo asymmetric divisions, producing a stem cell and a progenitor daughter cell. In many tissues, these stem cell progenitors further proliferate for several rounds, in a process known as transit amplification (TA) divisions, before the terminal differentiation (Lajtha, 1979; Rangel-Huerta & Maldonado, 2017). The stem cells are slow cycling cells which retain their mitotic potential for a long duration; whereas their transit-amplifying daughters proliferate relatively rapidly and have a limited mitotic potential (Cotsarelis et al, 1990; Potten & Loeffler, 1990; Rangel-Huerta & Maldonado, 2017).

Previous reports suggest that this difference in the stem cells and the TA cells could be partly attributed to distinct mechanisms of cell cycle regulation (Rangel-Huerta & Maldonado, 2017; Zaveri and Dhawan, 2018). A study in mouse pituitary gland showed that Cdk4, involved in the G1 cell cycle progression (Sherr, 1995), is selectively expressed by the TA cells, and its overexpression in stem cells leads to depletion of the stem cell pool, indicating a TA stage-specific role of Cdk4 (Macias et al., 2008). Moreover, p21 and p27 (cyclin-dependent kinase inhibitors) knockout studies in the mouse hematopoietic cells indicate that p21 specifically regulates stem cell cycle kinetics, whereas p27 regulates TA cell cycle kinetics (Cheng et al, 2000a, 2000b). Despite such differences, the division rates of stem cells and transit amplifying pool are often altered in a coordinated fashion by hormonal stimulation (Giraddi et al., 2015), tissue damage (Lehrer et al, 1998; Ichijo et al, 2017), and aging (Charruyer et al., 2009). A concerted analysis of the cell cycle regulation in stem cells and at different stages of subsequent TA is required to uncover the correlation and consequence of the stage-wise alterations of cell division rates on tissue growth and differentiation.

*Drosophila* spermatogenesis is a well-suited model system for assessing the impact of different cell intrinsic and extrinsic factors on the rates of GSC and TA divisions. The testis apex harbors ∼10 germline stem cells (GSCs) and ∼20 somatic cyst stem cells (CySCs). Asymmetric division of the GSCs produces the germline progenitor, gonialblasts (GBs), and that of CySCs produces the somatic cyst cells (SCCs). Each GB undergoes four rounds of symmetric TA divisions within an enclosure formed by two SCCs, forming cysts containing 2, 4, 8 and 16 interconnected germline cells. Subsequently, all the 16 germline cells within a cyst enter meiosis (Hardy, 1979; Fuller, 1998). The GSC division periods, estimated using the time-lapse imaging of intact testis preparations *ex vivo*, suggested that the proportions of symmetric and asymmetric rates of divisions are altered during regeneration (Sheng & Matunis, 2011). Also, periods of different cell cycle phases in GSCs are intrinsically regulated by Rho/ROK signaling and AuroraB kinase (Lenhart & DiNardo, 2015). Together, the evidence indicates that the cell cycle rates, governed by a combination of factors in stem cells, and at different stages of TA in *Drosophila* testis could be adaptable to the environment. However, the molecular specificities of the cell cycle regulation at the TA stages, and the impact of the variations in the cell cycle rates on the subsequent differentiation are still unclear.

Although multiple studies have modeled the dynamic changes in the stem cell behavior during tissue growth and homeostasis (Lei et al, 2014; Deasy et al, 2003; Hannezo et al, 2014), they are primarily focused on the stem cell behavior. Here, we describe a simplified mathematical analysis technique that utilizes the demographics of GSCs and cysts at different stages to arrive at probabilistic predictions of their lifespans at steady-state in adult testis. Numerical solutions of the equations using the observed distribution of live and dead cysts in fixed testis preparations suggested that halfway into the germline TA, the cell division periods are reduced by nearly 50%. This shortening of the cell division periods coincided with the altered persistence of G1 and G2 phases at the 4 and 8-cell stages. We then applied the mathematical model to the data obtained from tissue-specific RNAi and overexpression backgrounds to assess the roles of different cell cycle molecules involved in the G1-S and G2-M cell cycle transitions in the regulation of the lifespans of stem cell and TA stages. The analysis indicated that suppression of Cyclin E and CDK1 function in stem cells and early germline would extend the inter-division lifespans of GSCs and the TA stages without disrupting the subsequent spermatid differentiation. On the other hand, the loss of these functions after the second mitotic division arrested the TA at the 8-cell stage and affected subsequent differentiation in two distinct manners. Altogether, these results suggest that cell cycle regulatory proteins differentially influences the division rates of the GSCs and their progeny, and thereby, control subsequent tissue growth and differentiation.

## Results

### The persistence pattern of cell cycle phases in germline cysts change after the second mitosis

In wild-type testes, the SCCs and GSCs, and the germline cysts are arranged in a radial manner (Figure 1A and B). After each TA division, the resultant cyst is displaced further away from the stem cell niche, termed as the hub. The number of germ cells within a cyst, covered by the somatic cell membrane marked by Armadillo, identifies its stage (Figure 1C). Upon quantification of the number of cysts at each TA stage in testes from the wild-type and control genetic backgrounds, we found that the cyst numbers reduce significantly after the 2-cell stage (Figure 1D). To investigate whether germ cell death (GCD) at the 4-cell stage could cause the drop in the cyst number, we quantified GCD. Consistent with a previous estimate (Yang & Yamashita, 2015), we identified a few Lysotracker and Vasa-positive cysts at each TA stage (Figure S1B, C and Table S1). We did not find any sharp increase in cell death at the 4-cell stage. This observation indicated that increased GCD, by itself, cannot lead to such a bimodal cyst distribution. Therefore, we conjectured that the germline cyst might divide with an increasing rate during the second half of the TA.

**Figure 1:**
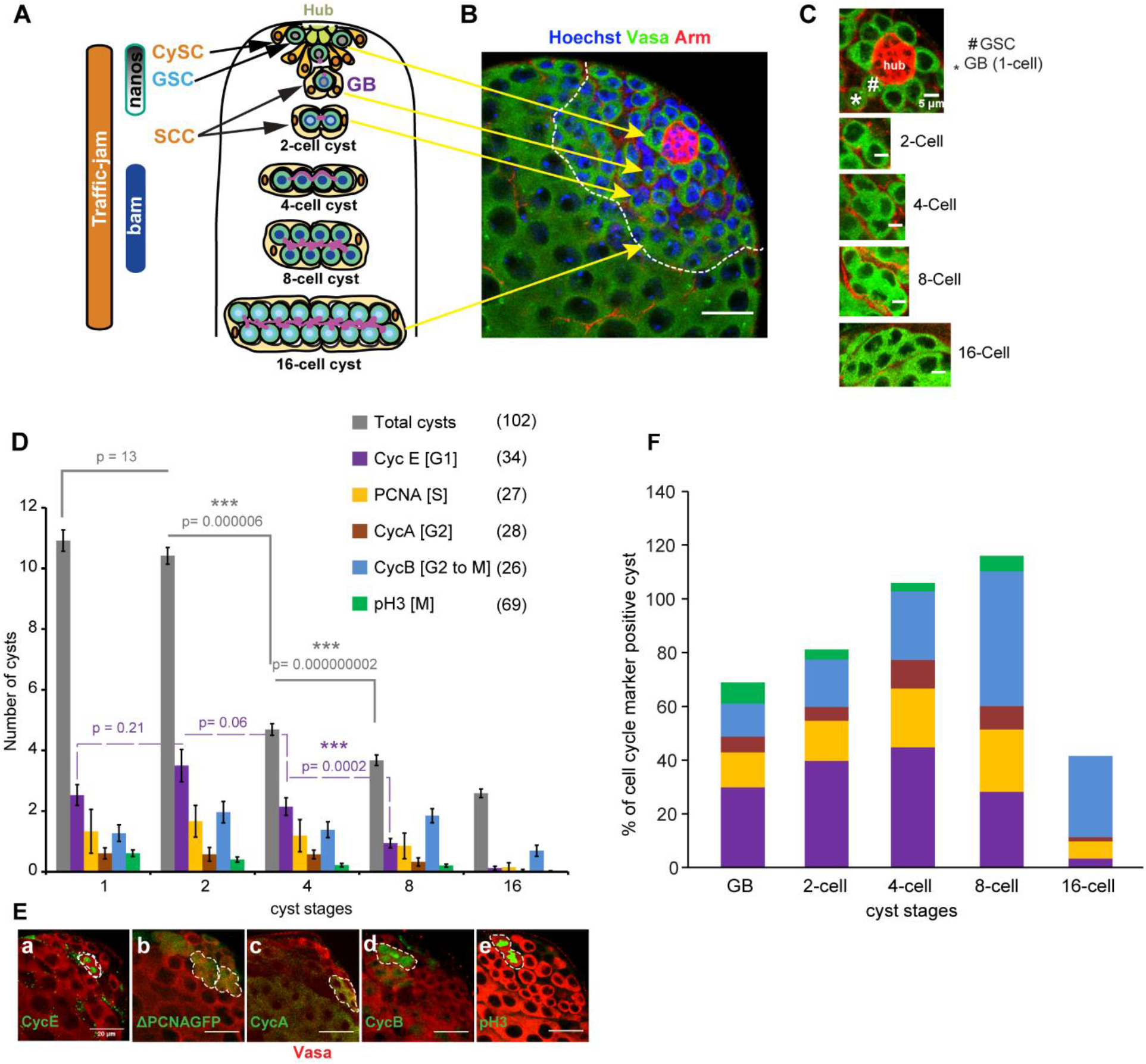
Stage-wise distribution of germline cysts and that of the cell cycle phases during the TA. A) Schematic illustrates the process of transit amplification during early spermatogenesis and expressions domains of the germ cell markers, nanos (nos) and bag-of-marbles (bam), and a somatic cell marker traffic-jam *in Drosophila*. Glossary: GSC – Germline Stem Cell, CySC – Cyst Stem cell, SCC – Somatic Cyst Cell, GB – Gonialblast, SG – Spermatogonia. B) The apical tip of wild-type (WT) testis stained with the Hoechst-dye (blue), anti-Armadillo (Red) and anti-Vasa (green). GSC, GB, 2-cell, and 16-cells are indicated by arrows. (scale bars ∼20μm). C) Enlarged images of the hub, GSCs, and cysts at different stages of TA in wild-type testis. (scale bars ∼5μm). D) Stage-wise distribution (average ± S.E.M) of germline cysts (grey bars) and cysts stained with various cell cycle markers (average ± S.E.M.) in the WT background. The pair-wise significance of difference was estimated using the Mann-Whitney-U test, and the p-values and sample numbers are indicated on the plot. E) WT testes stained with vasa (Red) and different cell cycle markers (green) indicate the distribution of cell cycle phases in the germline of cysts at different stages. Although it marked very few cysts in testis, the CycE (a), CycA (c), and PH3 (e) immunostaining and the expressions of ΔPCNA-GFP (b) and GFP-CycB protein trap (d) always marked all the germline cells enclosed within a cyst. (scale bars ∼20μm). F) Histogram shows relative frequency of cysts stained with different cell cycle phase markers at each stage of the TA. The ratios were calculated by dividing an average number of cell cycle positive cyst by the average number cyst at that stage (data from Figure.1).

To understand whether there is any correlation between the cell cycle phases and the rates of TA divisions, we scored the number and relative proportion of cysts marked for different cell cycle stage markers, spanning from G1 to M phases in wild-type testes (Figure 1D, E, F). A previous study in *Drosophila* ovary had reported a progressive shrinkage of the G2 phases after the second TA division, and an increase in the length of M and S phases (Hinnant et al. 2017). However, these results were based on Fly FUCCI reporter genes driven by *nosGAl4vp16 (nos>)*. *nos>* driver leads to a non-uniform expression of the downstream UAS transgene (Hinnant et al. 2017), which can skew the cell cycle phase distributions. Such false negative results were apparent in cases where the germ cells were positive for 5-ethynyl-2′-deoxyuridine (EdU) or Phospho-histone3 (pH3) but expressed no FUCCI marker genes (Hinnant et al. 2017). In contrast to their observations, we observed that at the 8-cell and 16-cell stage, the percentage of cysts marked by Cyclin E decreased, and the percentage of cysts expressing Cyclin B increased (Figure 1E and F). These results indicated that the cell cycle lengths of the TA stages, particularly the G1-S and G2-M transition periods, are likely to be altered after the second TA division.

### Mathematical analysis of stage-wise cyst distribution revealed a two-fold reduction in the lifespans of germline cysts after the second TA division

To estimate the inter-division lifespans of the germline cysts during the TA stages from these enumerations, we developed a mathematical model. We reasoned that in a closed system at steady state, the proportion of cysts of a given stage would be equal to the proportional lifespan of that stage, assuming that no cyst is lost due to death during the TA. We found that the relative distribution of cysts, calculated as the proportion of cysts of a particular stage with respect to total cysts, was invariant in the adult testis until 8-days after eclosion (Figure 2A), indicating that the rates of TA divisions remain in a steady state during this period. To account for the observed GCD (Figure S1B), we calculated the probability (*s*_*i*_) of a successful transition from one cyst stage to the next, as a function of the number of detected dead cysts (*D*_*i*_) and the time of their persistence (*A*) in the tissue, termed as the clearance time. Assuming that the number of dying cysts at the transition from stage *i* to stage *i* + 1 is *D*_*i*_, and expressing the quantity *s*_*i*_ in terms of the clearence time *A* and *D*_*i*_, we get the following equation.

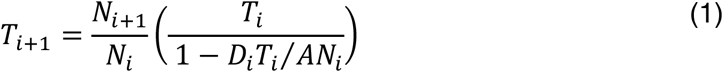

**Fig 2.**
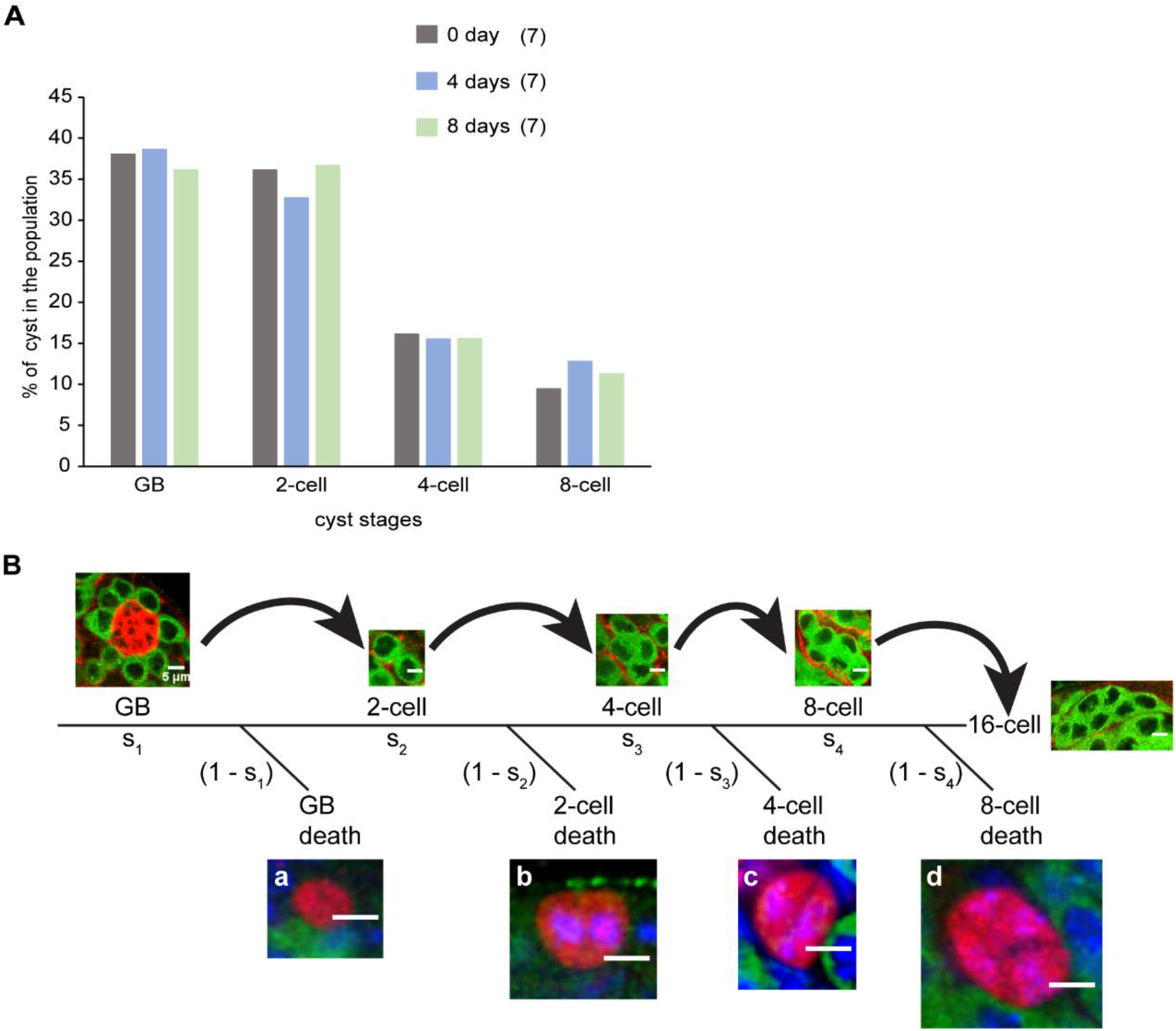
The cyst distribution is maintained at a steady state in adult testis at least until 8-days after eclosion. A) Histogram shows the relative stage-specific distribution profile of cysts (average) in wild-type (Canton S) adults aged for 0-, 4- and 8-days after emergence from the pupal case (eclosion) at 29°C. B) The schematic depicts the journey of a cyst from GB to a 16-cell cyst during the TA divisions with examples of GCD at each stage, e.g., GB (a), 2-cell (b), 4-cell (c) and 8-cell (d) marked by the lysotracker staining (Scale bars 5μm). The branches of the tree depict the probabilities of different outcomes. The time periods predicted for the first four TA divisions in wild-type testes have been indicated on the top.

In the above equation, *N*_*i*+1_ and *N*_*i*_ represent the average number of cysts observed at stage *i* + 1 and *i* respectively, quantified from confocal images of isolated testes from different genetic backgrounds. *T*_*i*+1_ and *T*_*i*_ are the lifetimes of stage *i* + 1 and *i* respectively (Refer to Supplemental Methods for a detailed reasoning). Inclusion of the possibility of death in our model converts a simple linear succession of stages into a branched system, with a terminal branch at each vertex (Figure 2B). Consequently, the simple linear ratios of numerical frequencies to periods used in a previous study for pupal ageing (Bainbridge and Bownes, 1981), now gets replaced by Möbius transformation. Such a transformation is non-linear, and it can lead to some unexpected behaviors near its singularity.

We calculated *D*_*i*_′*s* from the average number of dead cysts at each TA stage (Figure S1B). To calculate *A*, we needed to estimate the persistence times of dying cysts. Germ cell death marked by Lysotracker staining in *Drosophila* testis is known to occur in four sequential phases that are identified by the staining intensity and pattern (Chiang et al, 2017). Amongst these, phase-I to II transition is identifiable by an increase in the intensity of Lysotracker staining and the dye entry into the germ cell nucleus. Therefore, quantification of dead cysts was restricted to Phase-I stage, and “*A*” in Eq (1) was redefined as the Phase-I persistence time (Table S2). To estimate the average value of “*A*”, we collected time-lapse images of Lysotracker-stained testes, which revealed that all the TA stages exhibit a large variation in the Phase-I duration (Table S2 and Figure S2). Hence, we carried out a limited simulation assuming the values of “*A*” as 0, 1, 2, 3, 4 and 5 hours. Our results indicated that for the genotypes under investigation, this range of clearance time does not influence the time predictions to a considerable extent. Therefore, we assumed the Phase-I duration as 3 hours for all stages as it falls within the range of estimated values (Table S2).

### The duration of mitoses remains unaltered between GSCs and subsequent TA stages

In addition to the inputs of cyst numbers, frequency of death and persistence of dying cysts, the Eq.1 contained two more variables, viz, *T*_*i*_ and, *T*_*i*+1_. To begin the iterative determination of *T*_*i*_′s, we were required to estimate the lifespan of at least one stage from the GSCs and the four TA stages. We estimated the time period and frequency of the M-phase in GSCs, to calculate the lifespan of GSCs (see Supplementary Methods for further details). According to the time-lapse imaging analysis, the M-phase period is reported as 40 minutes to 1 hour (Sheng & Matunis, 2011). Since this was an indirect estimate, we measured the M-phase period of GSCs using time-lapse imaging *ex vivo*. The onset and termination of M-phase were monitored using Jupiter-GFP protein trap, which marks the astral microtubule and spindle body (Karpova et al., 2006), in *nosGal4vp16 UAS-Histone-RFP/Jupiter-GFP* background (Figure 3A). We considered the separation of centrosome-associated microtubules as the beginning of prophase (Lenhart and DiNardo, 2015; Siller et al., 2005), which is visible as bright fluorescent dots (arrowhead, Figure 3A-a). Subsequently, it separated and migrated to opposite ends of the nucleus (Figure 3A-b), and formed a spindle body characteristic of metaphase, with chromatin marked by His-RFP aligned at the cell equator (Figure 3A-c), then the spindle resolved through anaphase (Figure 3A-d) and telophase (Figure 3A-e). Following nuclear separation, the Jupiter-GFP localization persisted for a considerable duration marking the abscission period (Figure 3A-f and g). Finally, a visible increase in the nuclear size during the time-lapse series was identified as the end of telophase (Figure 3A-g). Accordingly, the M-phase period was estimated to take an average of 70 minutes for GSCs.

**Fig 3.**
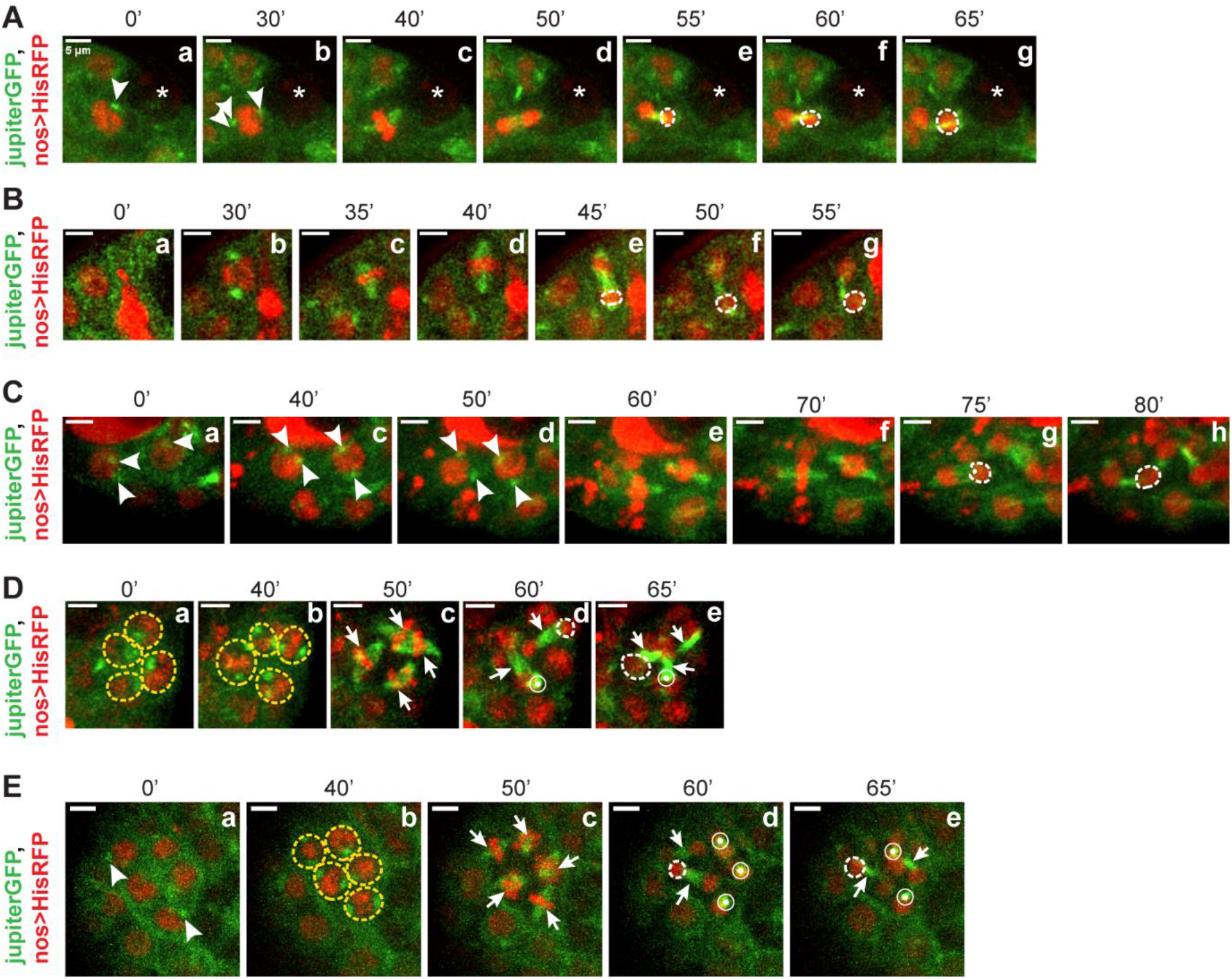
Time-lapse imaging of the GSC and TA cells division. Montage of a time-lapse movie of a GSC (A), GB (B), 2-cell (C), 4-cell (D) and 8-cell (E) (*JupiterGFP* (green), *nos>HisRFP* (red)), transitioning from prophase (a, b), metaphase (c, d), anaphase (e) and telophase (f). Arrowheads mark the position of the separated centrosomes. Arrows mark the metaphase plate and concentric circles mark the spindles perpendicular to the plane of the image. End of telophase is marked by an increase in the nuclear size marked by white dashed circles. Yellow dashed circles label the cells visible in the plane of imaging. Time intervals in minutes have been indicated in the top panels and scale bars ∼ 5μm.

The previous study in *Drosophila* ovary reported that the M-phase period varies significantly across TA stages based on the mitotic indices of different TA stages (Hinnant et al, 2017). Estimation of M-phase periods using time-lapsed imaging (Fig. 3B-F) indicated that M-phases of all the TA stages are similar to that of the GSCs (Figure 3B-E). These observations suggested that the time required to complete the M-phase remains invariant throughout the TA period. In line with our earlier observation, these estimations indicated that the cell cycle remodeling during TA period could occur during the G1 and G2 phases.

### The cyst lifespans reduce by nearly 2-folds after the second TA division in adult testis

We estimated the GSC lifespan in *nos>* background using the ratio of M-phase period to the GSC mitotic index (Eq.2, Supplemental Method, Table S3). The GSC lifespan was calculated to be 11.1 hours, which was marginally shorter than the time reported earlier in the same genetic background (Lenhart and DiNardo, 2015). This reduction was attributed to the rearing of stocks at 29°C. We then used Eq.1 to calculate the lifespans for cyst stages in the same background (Table 1). It indicated that the lifespans of GBs (*T*_1_, 1-cell stage, 13.6 hours) and 2-cell stage (*T*_2_, 12.3 hours) during the TA (Table 1) are comparable to that of the average interval between successive M-phases of a GSC (*T*_0_, 11.1 hours). In comparison, the life spans of 4-cell (*T*_3_, 6.9 hours) and 8-cell (*T*_4_, 5.8 hours) stages are progressively shortened (Table 1). In other words, the GBs and germline cells in 2-cell stage divide at the same rate as GSCs, whereas those in the 4 and 8-cell cysts divide at ∼50% faster rates. These results indicated that along with the cell cycle phase remodeling, duration of the entire cell cycle is altered after the second TA division.

**Table 1:**
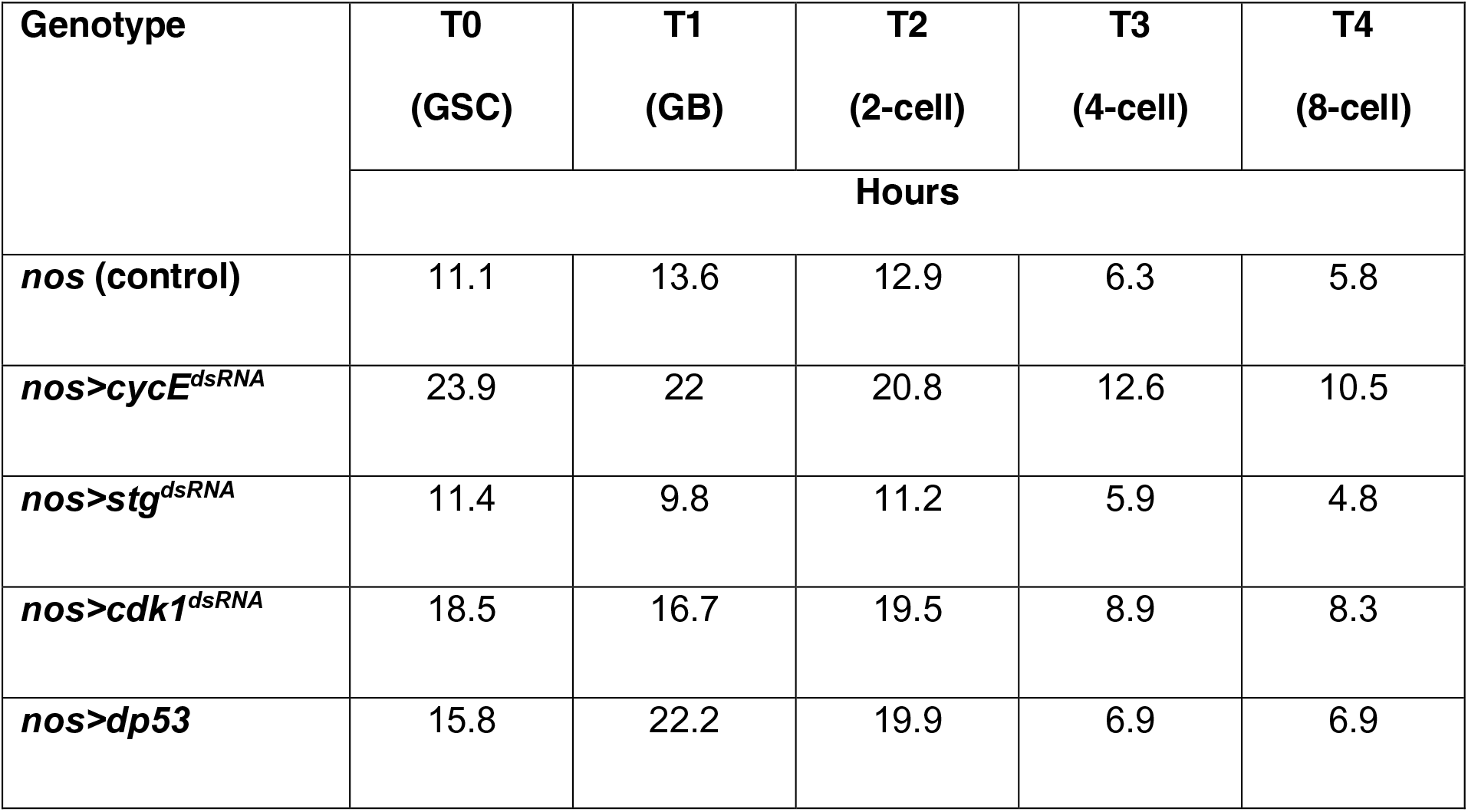
Estimated persistence time of different cyst stages during the TA in different genetic backgrounds.

| Genotype | T0<br>(GSC) | T1<br>(GB) | T2<br>(2-cell) | T3<br>(4-cell) | T4<br>(8-cell) |
| --- | --- | --- | --- | --- | --- |
|  | Hours |  |  |  |  |
| ***nos* (control)** | 11.1 | 13.6 | 12.9 | 6.3 | 5.8 |
| ***nos>cycE<sup>dsRNA</sup>*** | 23.9 | 22 | 20.8 | 12.6 | 10.5 |
| ***nos>stg<sup>dsRNA</sup>*** | 11.4 | 9.8 | 11.2 | 5.9 | 4.8 |
| ***nos>cdk1<sup>dsRNA</sup>*** | 18.5 | 16.7 | 19.5 | 8.9 | 8.3 |
| ***nos>dp53*** | 15.8 | 22.2 | 19.9 | 6.9 | 6.9 |

### Perturbations of the cell cycle checkpoint control in the GSCs and early germline alter the TA pool

To quantify the roles of the cell cycle regulatory proteins in determining the lifespans of GSCs and the TA pool, we perturbed the functions of two different cell cycle regulators – Cyclin E and CDK1, in GSCs and early germline cysts (Figure 4). We used *nos>*, which expresses in GSCs and early germline cells (Figure 4B-a), to drive the expression of different transgenes. We modulated the length of the G1 phase by RNAi of cyclin E (cycE) and dacapo (dap), and overexpression of a stable form of CycE (CycE^ΔEPEST^) protein, respectively (Figure 4B-c,d,e). Similarly, we perturbed the length of the G2 phase in GSCs and early germline using RNAi of string (stg), and cdk1, as well as overexpression of *wee1*, respectively (Figure 4B-f,g,h). CycE drives the G1-S transition (Duronio et al., 1998; Duronio and O’Farrell, 1994; Follette and O’Farrell, 1997; Lee and Orr-Weaver, 2004). Dap is a member of the p21/p27 family of Cdk inhibitors which inhibits CycE/Cdk2 activity (De Nooij et al., 1996). *dap* mutant embryos undergo an extra division cycle, and ectopic overexpression of dap arrests the cells at G1 (Lane et al., 1996). Cdk1, the Cdc2 homolog in *Drosophila,* is required for the G2-M transition (Clegg et al., 1993; Stern et al., 1993). Wee1 is an inhibitory kinase which inhibits Cdk1 by phosphorylating a tyrosine residue (Campbell et al., 1995), and *stg* codes for a Cdc25 phosphatase which removes this inhibitory phosphorylation on Cdk1, facilitating the G2-M transition (Edgar and O’Farrell, 1989; Gautier et al., 1991; Kumagai and Dunphy, 1991).

**Fig 4.**
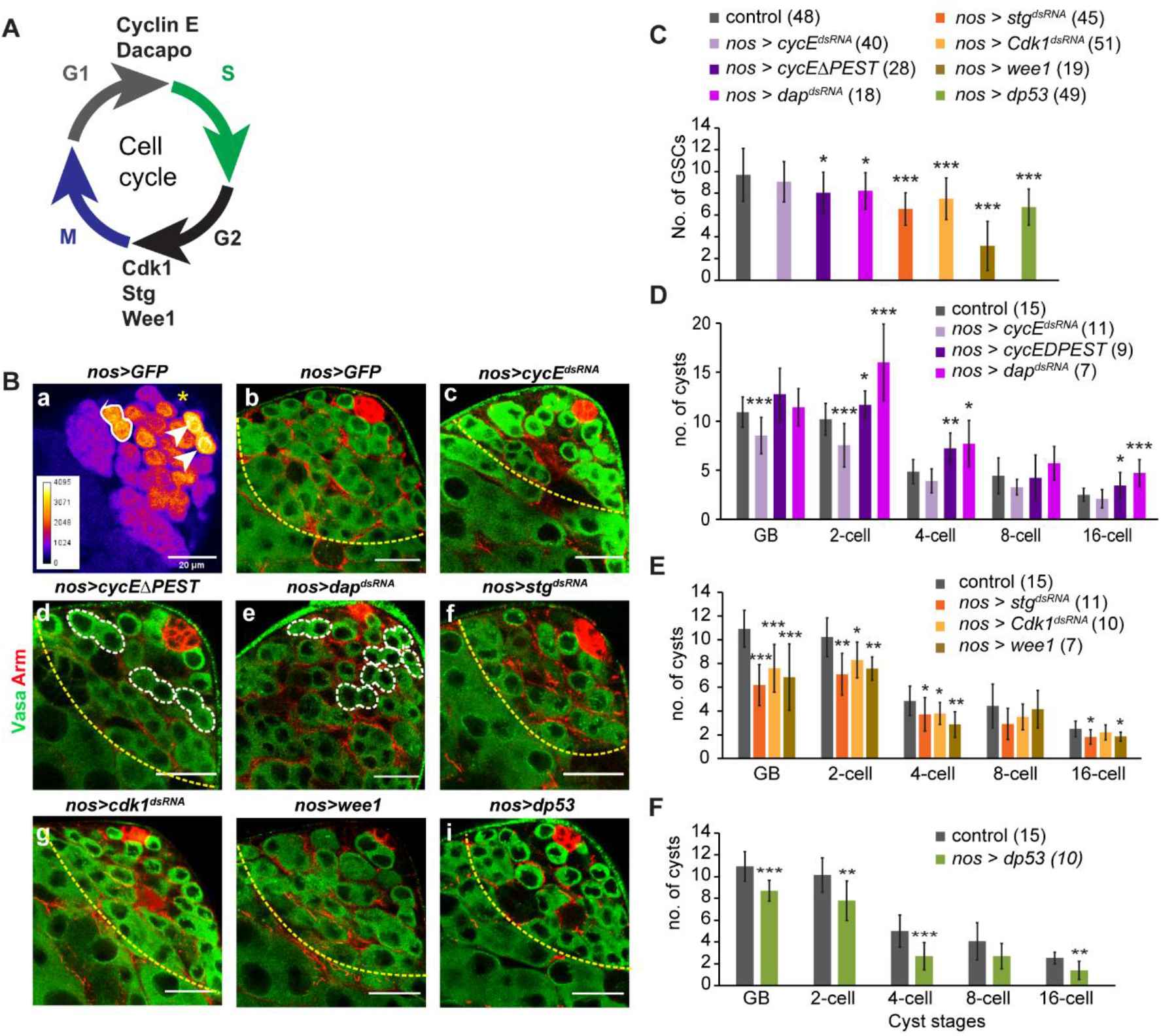
Stage-wise distribution of cysts after perturbations of the cell cycle checkpoints in the GSC and subsequent early germline progeny cells. A) The schematic depicts a typical cell cycle in the germline with G1, S, G2 and M phase with molecules under study categorized as the molecules involved in G1 to S transition (cyclin E and Rbf1) and G2 to M transition (stg, Cdk1, Wee1). B) Apical tip of an adult testis from *nos>EGFP* background showing the expression of UAS-EGFP (FIRE LUT) (a). Testes from *nos>EGFP* (b), *nos>cycE^dsRNA^*(c), *nos>cycE^ΔPEST^* (d), *nos>dap^dsRNA^* (e), *nos>stg^dsRNA^* (f), *nos>Cdk1^dsRNA^*(g), *nos>wee1* (h) and *dp53* (i) flies, stained with the Hoechst dye (blue), anti-Arm (Red), Lysotracker (red) and anti-Vasa (green), show shrunken apical tip with fewer cysts as compared to control. (scale bars ∼20μm). Germline TA region is marked with a yellow dotted line. C) The histograms depict the number of GSCs (average ± S.D.) in *nos>eGFP, nos>cycE^dsRNA^, nos>cycE^ΔPEST^, nos>dap^dsRNA^*, *nos>stg^dsRNA^, nos>Cdk1^dsRNA^, nos>wee1* (D), and *nos>dp53* backgrounds. D-F) The histograms depict the stage-wise distribution (average ± S.D.) of cysts in *nos>EGFP, nos>cycE^dsRNA^, nos>cycE^ΔPEST^, nos>dap^dsRNA^* (B), *nos>stg^dsRNA^, nos>Cdk1^dsRNA^, nos>wee1* (D), and *nos>dp53* (E) backgrounds. The pair-wise significance of difference was estimated using the Mann-Whitney-U test (p-values are * <0.05, **<0.01, ***<0.001) and the sample numbers are as indicated on the histogram panels.

In *Drosophila* ovary, loss of CycE reduces the number of GSCs (Ables and Drummond-Barbosa, 2013). We observed that the number of GSCs remained unaltered in *nos>cycE^dsRNA^*testes, and they were significantly reduced in *nos>cycE^ΔPEST^*and *nos>dap^dsRNA^* testes, respectively (Figure 4C). Together these results suggested that elevated level or activity of CycE is detrimental to the maintenance of GSC pool in *Drosophila* testis. A previous study has shown that Stg is also required for maintenance of GSC number (Inaba et al, 2011). Consistent with this report, we also found significant reductions in the GSC numbers in *nos>stg^dsRNA^, nos>cdk1^dsRNA^* and *nos>wee1* testes, respectively (Figure 4C), suggesting that a timely exit from G2 might be necessary for stem cell maintenance.

CycE RNAi in GSCs and early germline cells in *nos>cycE^dsRNA^*testes also led to a significant reduction in the average number of cysts in 1-4 cell stages, but the number of 8-cell cysts was nearly the same as that of control (Figure 4D). In comparison, the expressions of *nos>cycE^ΔPEST^*, and *nos>dap^dsRNA^*, respectively, led to a significant increase in the average number of 2- and 4-cell cysts (Figure 4D). Since the highest levels of nos>GFP expression was observed in GSCs, and GBs, we conjectured that a selective acceleration of GSC and GB (1-cell) divisions could cause such an accumulation. The opposing effects observed due to the *nos>cycE^dsRNA^* expression, *vis-a-vis* that of the *nos>cycE^ΔPEST^*, and *nos>dap^dsRNA^* overexpression, corroborated that the CycE levels play a significant role during the TA stages. Hence, these results indicated that modulating the length of the G1 phase in the GSCs and the early germline could affect the TA divisions during the initial stage. A similar and more persistent loss was observed in *nos>cdk1^dsRNA^*, *nos>stg^dsRNA^*and *nos>wee1*, testes, which contained significantly reduced numbers of 1-4 cell stage cysts (Figure 4E). These observations further indicated that perturbations of the G2 exit in GSCs and the early germline cells could also affect the subsequent TA divisions in the early stages (Figure 4F). Together, these observations indicated that G1 and G2 modulations in the GSCs and the early germline stages could modify the distribution profile and overall growth of the germline pool in adult testis.

Overexpression of *Drosophila p53*, a DNA damage checkpoint protein known to induce apoptosis upon DNA damage (Fan et al., 2010; Ollmann et al., 2000), via *nos>dp53* also depleted the number of cysts at all the TA stages (Figure 4D). Hence, it suggested that the ectopic expression of p53 could also influence the cell division in the GSC and TA stages.

Although these quantifications indicated that the cell cycle regulatory molecules regulate the divisions of GSCs and TA stages, it could not provide a clear understanding of their influence on the stage-specific division rates of the germline pool. Also, one could not estimate the actual alterations in the inter-division lifespans due to these perturbations from the data alone.

### Suppression of the cell cycle controls in GSCs, and early germline differentially slowed down the rates of GSC and TA divisions

To further understand the impact of the cell cycle perturbations on the rates of GSC and TA cell divisions, we estimated the stage-wise distribution of mitotically active cysts immunostained with pH3 antibody in the mutant testes. In *Drosophila* ovary, *cycE* homozygous mutant GSCs and cysts did not incorporate EdU indicating a G1-S block (Ables and Drummond-Barbosa, 2013). Consistent with this result, we also observed a significant reduction of the average number of pH3-stained GSCs, as well as the mitotic index, in the *nos>cycE^dsRNA^* testis (Figure 5A-b and B). stg RNAi is also reported to reduce the mitotic index of GSCs, calculated as the average number of pH3-positive GSCs, in *Drosophila* testis (Inaba et al, 2011). This method of the mitotic index calculation used by Inaba et al, 2011 does not consider the total GSC pool in a given testis, and therefore, it does not normalize for variation in the GSC number across different genotypes. According to our estimates, both the average number of GSCs and the pH3-positive GSCs were reduced in *nos>stg^dsRNA^* testes (Figure 5B). Hence, the mitotic index was not significantly altered (Table S3). Similar results were obtained from the cdk1 RNAi, and dp53 overexpression backgrounds (Table S3). Together, these results indicated that loss in the GSC pool in the cdk1 and stg RNAi backgrounds might not indicate alterations in cell cycle periods.

**Fig 5.**
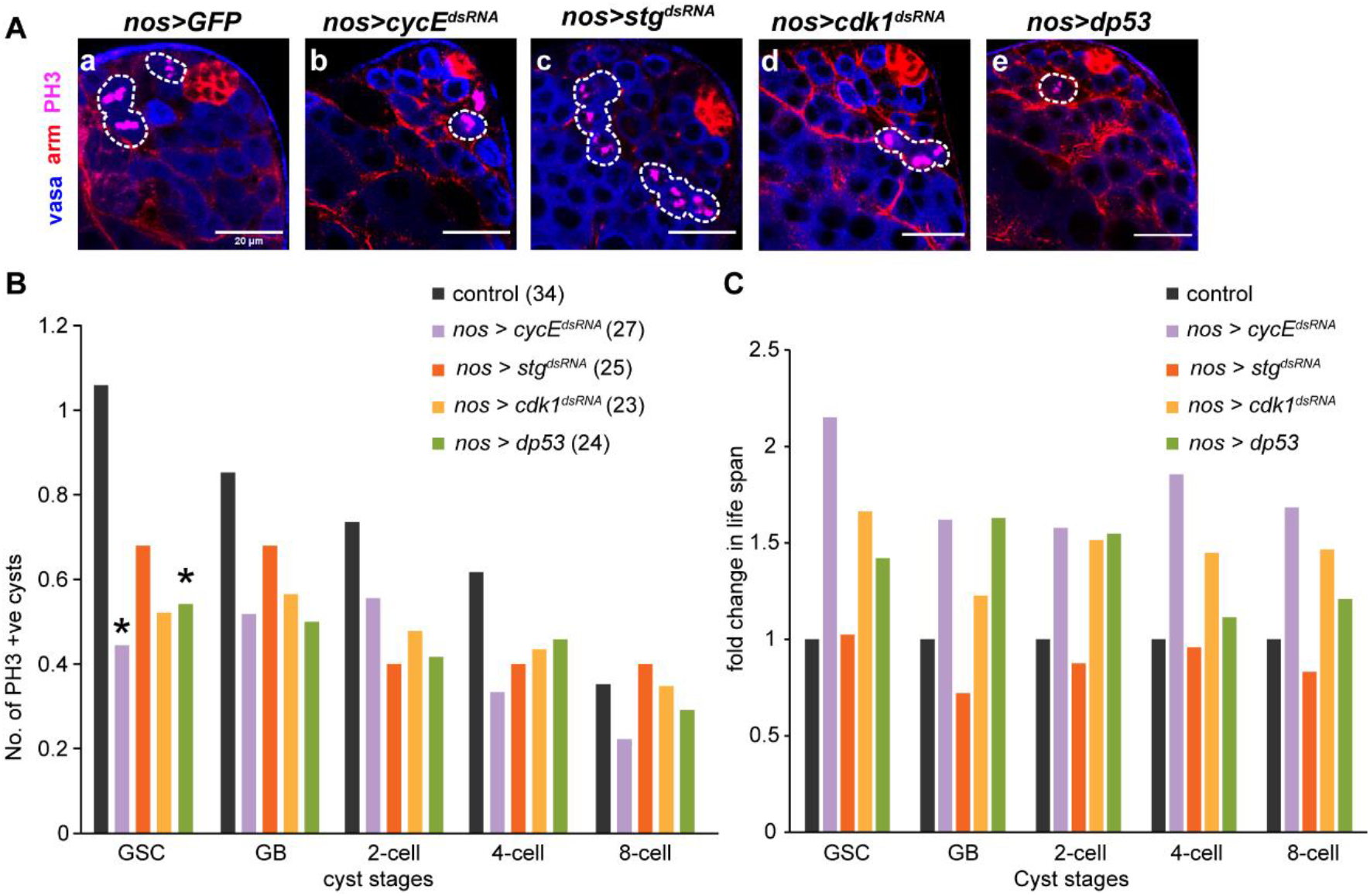
Stage-specific life-span of the germline cysts estimated after the germline-specific RNAi of the cell cycle regulatory genes and overexpression of dp53, respectively. A) Testes from *nos>EGFP* (a), *nos>cycE^dsRNA^*(b), *nos>stg^dsRNA^* (c), *nos>Cdk1^dsRNA^* (d) and *nos>dp53* (e) flies, stained with anti-Vasa (blue), anti-Arm (Red), and phospho-histone3 (PH3 in magenta). (scale bars ∼20μm). PH3 positive stages are marked with dotted outline. B) The histograms depict the stage-wise distribution (averages) of PH3-positive cysts in the *cycE*, *stg* and *Cdk1* RNAi, and dp53 overexpression backgrounds. The pair-wise significance of difference was estimated using the Mann-Whitney-U test (p-values are * <0.05, **<0.01, ***<0.001) and the sample numbers are as indicated on the histogram panels. C) The histograms depict the time periods estimated by applying equation-1 to data obtained from *nos>EGFP, nos>cycE^dsRNA^*, *nos>stg^dsRNA^*, *nos>Cdk1^dsRNA^* and *nos>dp53*.

Further, to analyze the effects of cell cycle perturbations on the inter-division lifespans of GSC and TA stages, we applied the mathematical equation (Eq.1) to the data from select candidates of our screen, *viz*, the *cycE, cdk1,* and *stg RNAi* backgrounds, and that of *dp53* overexpression. The output suggested that the lifespans of GSCs and subsequent TA stages are enhanced up to 2-folds in the *nos>cycE^dsRNA^* background (Figure 5C), indicating that CycE plays a critical role in shaping the cell division rates of both GSCs and TA stages (Table 1). RNAi of cdk1 enhanced the lifespans of GSCs and TA stages to a comparatively lesser extent, which could be attributed to the efficacy of the dsRNA construct in these cells. stg knockdown, however, did not alter the lifespans of GSCs and TA stages, even though Stg has been implicated in GSC maintenance (Inaba et al, 2011). Altogether, these predictions suggested that the cell cycle regulatory genes intrinsically regulate the TA stages, and further validated the mathematical model.

A previous report has suggested that in *Drosophila* male and female germline, p53 activity remains limited to GSCs upon radiation-induced DNA damage (Wylie et al., 2014). Thus, the enhanced lifespans caused by dp53 overexpression as indicated by this study could be an artificially induced, ectopic overexpression defect. These results highlighted the effectiveness of the model in extracting the underlying differences in the rates of GSC and TA divisions which were not often apparent from the cyst distribution profiles.

### CycE and Cdk1 RNAi in 4 and 8-cell stage arrested the TA at the 8-cell stage and induced premature meiosis

Next, to study the regulation of the cell cycle phases after the transition to the rapid TA divisions, we expressed the *cycE^dsRNA^, stg^dsRNA^*, *cdk1^dsRNA^, wee1* (Figure 6) and *dp53* transgenes (Figure S2) using 2x*bamGal4*. Since we wanted to examine the effect of these perturbations on the spermatocyte population, we counted all cysts in the testis apex within ∼135μm from the hub. This area consists of the GSCs, the TA stages, and the spermatocyte pool including the S3-spermatocyte stage (Gupta and Ray, 2017). The *bam>cycE^dsRNA^* expression led to a marginal increase in the GSC and GB pool, and substantial accumulation of the 8- and 16-cell cysts (Figure 6A-c, B), indicating that CycE controls the G1-S transition at the 8-cell stage. Since Stg expression is limited to the stem cells (Inaba et al. 2011), the expression of *stg^dsRNA^* transgene in 4-16 cell cysts did not alter the cyst distributions (Figure 6A-b, B). In *bam>cdk1^dsRNA^*testes, the number of 4-cell cysts increased by ∼2-fold, these testes were filled with 8-cell cysts and had no 16-cell cysts (Figure 6A-d, B). A similar phenotype was observed with Wee1 overexpression (Figure 6A-e, B). Together, these two results suggested that Cdk1 activity is essential for the G2-M transition at the 8-cell stage.

**Fig 6.**
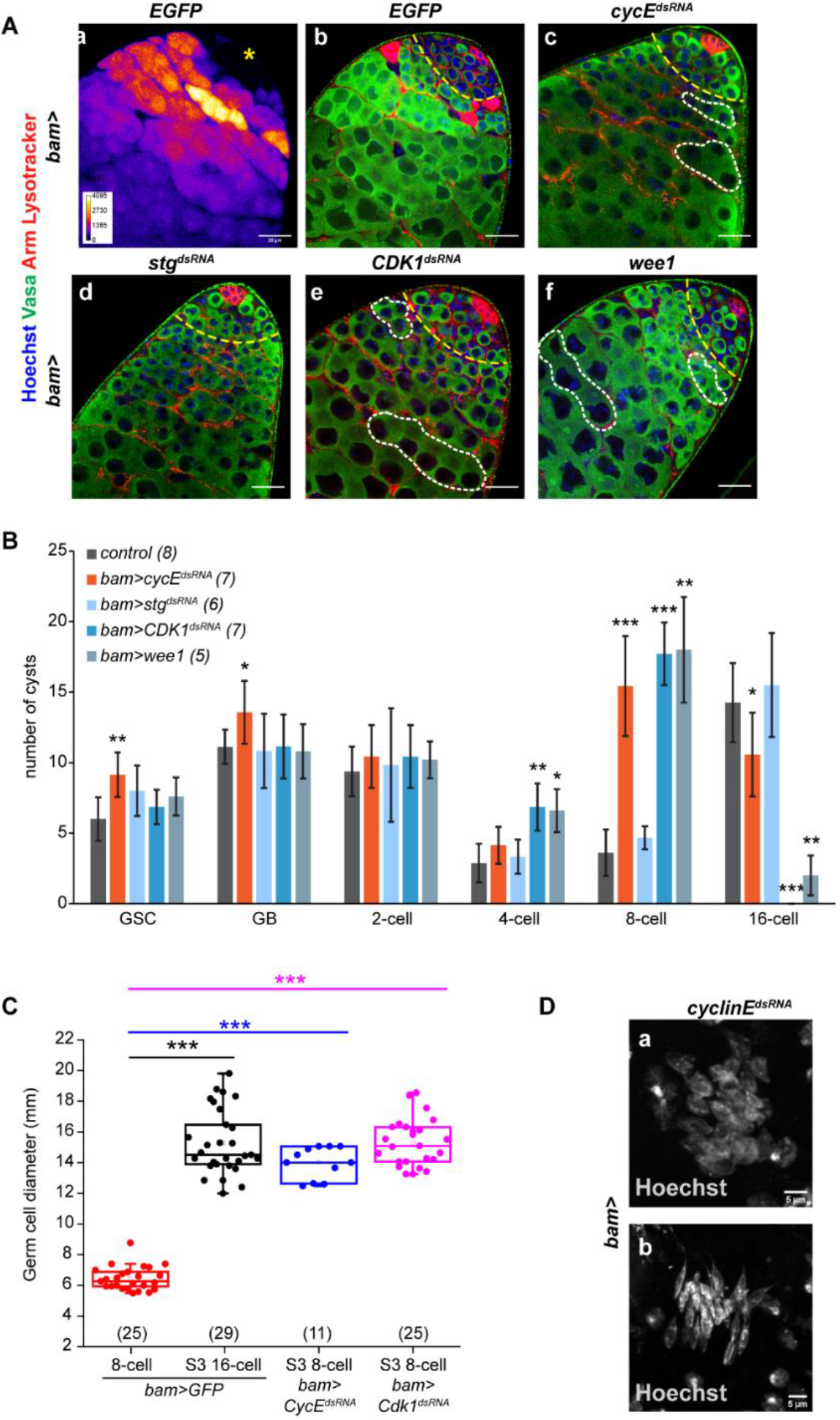
Stage-wise distribution of cysts in the backgrounds of late germline-specific RNAi of CycE, cdc25/string (stg), and Cdk1, respectively, and wee1 overexpression. A) Apical tip of an adult testis from *bam>EGFP* background showing the expression of UAS-EGFP (FIRE LUT). Testes from *bam>EGFP* (a), *bam>cycE^dsRNA^*(b), *bam>stg^dsRNA^* (c), *bam>Cdk1^dsRNA^* (d) and *bam>wee1* (e) flies, stained with the Hoechst dye (blue), anti-Arm (Red), Lysotracker (red) and anti-Vasa (green), show shrunken apical tip with fewer cysts as compared to control (scale bars ∼20μm). Dotted outlines indicate some of the abnormally accumulated 8-cell cysts. B) The histograms depict the stage-wise distribution (averages ± S.D.) of cysts in the CycE, stg, and Cdk1 RNAi, respectively, and in the wee1 overexpression, backgrounds. The pair-wise significance of difference was estimated using the Mann-Whitney-U test (p-values are * <0.05, **<0.01, ***<0.001) and the sample numbers are as indicated on the histogram panels. C) The nuclear diameters of the 8-cell spermatogonial, and 16-cell, S3-stage spermatocyte cysts from the *bam>EGFP*, *bam>cycE^dsRNA^*and *bam>Cdk1^dsRNA^*, backgrounds. The pair-wise significance of difference was estimated using the Mann-Whitney-U test (p-values are * <0.05, **<0.01, ***<0.001) and the sample numbers are as indicated on the graph. D) Examples of a post-meiotic cyst with 32 nuclei (a) and a bundle of elongated nuclei of 32 spermatids in the *bam>cycE^dsRNA^*testis (scale bars ∼5μm).

*bamGal4>dp53* overexpression caused widespread germ cell death (Figure S3A, arrows). Also, it contained cysts with more than 32 cells (Figure S3A, arrowhead) with a compact chromatin morphology, similar to that of the spermatogonia (Figure S3Bb, arrowhead). Due to punctate and irregular Armadillo immunostaining (Figure S3A), we could not estimate the cyst distribution profile in these testes; but the GSC counts were not significantly different than that of the control (Figure S3C). These results suggested that dp53 overexpression in the later stages of TA could be detrimental for meiotic entry and progression.

To investigate whether the cysts arrested at the 8-cell stage in the *bam>cycE^dsRNA^* and *bam>cdk1^dsRNA^*backgrounds could undergo meiosis, we measured the nuclear diameters of germline cells at 8-cell and S3 spermatocyte stages in the respective RNAi backgrounds. We found that the nuclei of the arrested 8-cell cysts in *bam>cycE^dsRNA^* and *bam>cdk1^dsRNA^*testes had similar sizes as that of the S3-stage spermatocytes in control testes (Figure 6C). These results suggested that the cell cycle arrest at the 8-cell stage caused by the knockdown of CycE and Cdk1 could lead to premature entry into meiosis. We also observed post-meiotic 32-cell cysts (Figure 6D) and elongated sperm tails (Figure S4B) in *bam>cycE^dsRNA^* testes. In contrast, we did not find any post-meiotic 32-cell cysts in both *bam>cdk1^dsRNA^* and *bam>wee1* testes. Together, these results indicated that the G1-S arrest due to the CycE RNAi at the 8-cell stage could induce premature meiosis and subsequent differentiation. These results also suggested that in addition to its role in late TA stages, Cdk1 also plays a critical role in regulating meiosis, which is consistent with a previous report that implicated *cdc2 (cdk1)* in meiosis (Sigrist et al., 1995).

## Discussion

### Progeny of the germline stem cells transitions to transit amplifying mode after the second TA division

Adult stem cells are maintained in a quiescent phase (G_0_) for extended durations with an occasional induction of cell cycle whereas transit amplifying cells cycle rapidly without the intervening G_0_ phase (Cheung & Rando, 2013; Lajtha, 1979). Here, we show that the progeny of the GSC divide at the same rate as the GSCs for two subsequent mitotic divisions in *Drosophila* testis. Following which, the TA divisions accelerate by nearly 2-folds. We found that the total proportions of GBs and 2-cell cysts marked by different cell-cycle-phase reporters relative to the total number of cysts at each of these two stages are less than 100%. In comparison, the sum of the proportions of cysts undergoing active cell-cycle in 4 and 8-cell stages exceeds 100%. The lifespan estimates also indicated that GSCs, GBs, and 2-cell cysts divide at a comparable rate whereas the germline cells in the 4- and 8-cell cysts divide at ∼2-fold faster rate. Therefore, similar to GSCs, the GBs and spermatogonial cells in 2-cell cysts could go through a quiescent (G_0_) phase in between active cell divisions, indicating a change in the cell cycle regimen after the second mitotic division. Although a similar cell cycle remodeling is reported at the 4/8 cell stage during the TA in *Drosophila* ovary (Hinnat et al., 2017), the conclusions were qualitative and did not comment about the length of the cell cycle. The mathematical modeling analysis used in this study further advanced the idea by providing lifespan estimates of these stages.

Incidentally, this transition coincides with the onset of Bam expression (Insco et al., 2009). Bam threshold plays a critical role in arresting the TA before the meiotic transition (Insco et al., 2009). *In vitro* and *in vivo* analysis in a previous study has suggested that Bam could stabilize Cyclin A (Ji et al., 2017). In *Drosophila*, ectopic Cyclin A expression has been shown to induce S phase entry in multiple tissues and stabilization of Cyclin A has been shown to arrest the cells in metaphase (Follette and O’Farrell, 1997; Sprenger et al., 1997; Thomas et al., 1994). We hypothesize that in the case of germline TA in the testis, the onset of Bam could play a role in remodeling the cell cycle by the stabilization of Cyclin A leading to shortening of the G1 phase and extension of the G2 phase.

### Regulations of the G1-S and G2-M transitions at the 8-cell stage differentially influences subsequent differentiation

In proliferating cells, the G1-S and G2-M transitions play pivotal roles in determining the cell division rates. CycE and associated proteins drive the cells through the G1-S checkpoint whereas CDK1 drives the G2-M transition. We found that these two regulators have differential roles in the early and late TA stages. RNAi of CycE in GSCs and early germline slowed the division rates of GSC and the subsequent TA divisions by ∼2-folds indicating that CycE predominantly regulates the divisions of both GSCs and TA cells. The overexpression of stable CycE^ΔPEST^ and dap RNAi in these cells also accumulated the intermediate TA stages, suggesting that regulating the CycE activity in the early germline would maintain their lifespans.

Similarly, we found that the CDK1 also plays a significant role in managing the lifespans of GSCs and TA stages. This result further indicated that along with CycE, Cdk1 also play an essential role in this process. Contrary to the earlier report which suggested that loss of Cdc25/Stg decreases the mitotic potential of GSCs (Inaba et al. 2011), we did not find any noticeable change in the lifespans of GSCs or the TA stages, although the GSC number was reduced in the stg RNAi background. The GSC pool is maintained by a combination of symmetric and asymmetric divisions, as well as back recruitment form the spermatogonial TA pool. Therefore, Stg could have a cell cycle-independent role in the maintenance of the GSC pool. Dp53 is expressed in the early spermatogonial cells (Napoletano et al., 2017) and is activated in response to radiation-induced DNA damage (Wylie et al, 2014). We found that dp53 overexpression could prolong the lifespans of GSCs and TA early stages.

RNAi of cycE and cdk1 at the late TA stages arrested the mitoses at the 8-cell stage, indicating that these two cell cycle phase regulators are essential for the progression of the final TA division. We further noted that only in the case of CycE RNAi some of the 8-cell cysts could complete meiosis and partly differentiate as elongated spermatids. Whereas in the *bam>cdk1 RNAi* and *bam>wee1* testes, the TA was arrested at the eight-cell stage, suggesting that the G2-M checkpoint is also critical for meiosis progression. This conclusion is consistent with the earlier studies, which reported that conditional loss of *cdc2* (Cdk1) could affect the meiotic progression in *Drosophila* testis (Sigrist et al., 1995). Loss of the dp53 gene was reported to cause tissue hyperplasia and an accumulation of the early spermatocytes which could indicate an accelerated transition from mitotic to the meiotic stage (Napoletano et al, 2017). We showed that *dp53* overexpression in the late stages of the TA induced cell death and blocked entry into meiosis. Together these results suggest that p53 activity and expression levels might be inhibited in primary spermatocytes to facilitate entry into meiosis.

### Mathematical modeling of the enumerated demographics elicited differential rates of the germline stem cells, early and the late TA divisions in *Drosophila* testis

Tissues adapt to the environmental changes such as nutrient depletion, hormonal stimulation, wound healing, *etc*. and transition between different states of homeostasis (Yang & Yamashita, 2015; Giraddi et al, 2015; Chiang et al, 2017; Lehrer et al, 1998). The quantitative data interpretation in this area has been limited to the stem cells (Roth et al, 2012; Rodgers et al, 2014; Seidel & Kimble, 2015; Cotsarelis et al, 1990; Inaba et al, 2011; Ojeh et al, 2015). Adult stem cells and the TA systems in various tissues often adapt by establishing a new steady-state under different growth conditions influenced by environmental and physiological changes (Yang & Yamashita, 2015; Giraddi et al, 2015; Chiang et al, 2017; Lehrer et al, 1998). The impact of the altered rates of stem cell divisions on subsequent TA has not been so well documented. A recent report utilized dual labeling method using BrdU incorporation to estimate the rates of cell division at steady state in mouse neocortex (Harris et al 2018). Although this method appeared simple, BrdU incorporation has been shown to induce toxicity (Levkoff et al., 2008; Taupin, 2007), which might skew the results.

Often the transit amplifying pool is considered as a homogenous proliferative population. The gene expression analysis in *Drosophila* ovary and testis, however, indicated a certain degree of genetic differentiation during the TA (Insco et al., 2009; Li et al., 2009). In *Drosophila* testis, the GSCs are easily identifiable due to the particular floral arrangement around the hub, whereas the TA cysts are more abundant and tightly packed with no specific spatial marker for identification in live tissue. Also, germline cells in a cyst are interconnected by ring canals (Fuller, 1993), which allow the intercellular exchange of proteins (Airoldi et al., 2011; McLean and Cooley, 2013), making it difficult to distinguish the GSC clones from TA cyst clones using the clone-based methods (Luo, 2007; Evans et al, 2009). Here, we showed that the mathematical modeling of the demographic distribution obtained from fixed tissue data could offer an alternative estimate of the rates.

Segregation of the germline progeny in the somatic-cell encapsulated cysts in testis helped to discriminate and enumerate the stage-specific distribution during the TA. Since all the TA cysts remain confined at the testis apex and the cyst distribution remains in a steady state during the early adult life, we could propose a probabilistic argument that the proportion of cysts at each stage would be equal to the proportional lifetimes if there is no loss due to germ cell death. We then incorporated the GCD and clearance time in the model to obtain the equation for prediction of the lifetimes of stem cell and TA stages. This technique allowed us to calculate the lifespans of different cell types within the tissue by enumerating the steady-state distribution in fixed preparations. The method demonstrated that one could estimate the average lifetimes of different demographic stages in a steady state system using simple data such as the progeny number, cell death, and mitotic index. Analysis of data obtained from control and genetically manipulated backgrounds using this technique yielded a clear distinction between the rates of the GSCs and stage-specific TA divisions.

This method can be easily used to obtain quantitative distinctions of the TA rates in the germline in different genetic backgrounds and those altered by environmental perturbations. The analysis of mutant data using this method helped to distinguish between similar results and suggested that the lifespans predictions are not always obvious from the demographic distribution. One of the limitations of this method is the requirement of stage-wise discrimination of cellular lineage to identify cells at different stages of transit amplification within a tissue. We envisage that one could adapt the formula by careful calibration of the dye tracing data in the progeny of other stem cells such as the type II neuroblasts in *Drosophila* (Boone & Doe, 2008; Homem & Knoblich, 2012), skin epidermal stem cells in *Zebrafish* (Guzman et al., 2013), mouse corneal epithelial stem cells (Kawasaki et al., 2006), and mammalian intestinal stem cells (van der Flier and Clevers, 2009).

In conclusion, our study shows that the germline divisions during *Drosophila* spermatogenesis speed up halfway through the TA. Extension of the G1 or G2 phases could slow down the cell cycle in the early germline stages; however, it arrested the divisions in later stages, indicating the differential impacts of the cell cycle regulatory proteins in maintaining the divisions of stem cells and the TA stages. We also showed that cell cycle arrest at the final TA division promotes premature entry into meiosis. Here, we also described a mathematical model which could predict the periods of the cell divisions using demographic data obtained from fixed tissue analysis. This method can be used to study the effect of various extrinsic and intrinsic factors on rates of stem cells and their progeny divisions, making it an attractive model for stem cell biologists.

## Materials and methods

### Drosophila stocks and culture condition

All stocks and crosses were maintained on standard *Drosophila* medium at 25°C unless mentioned otherwise. The flies were reared for four days at 29°C before dissection and fixation as described before (Joti et al, 2011). A detailed stock list is mentioned in Table S8.

### Whole-mount immunofluorescence-staining

Testes from a four-day-old male were dissected in Phosphate buffer-saline (PBS) and fixed in 4% paraformaldehyde for 20 to 30 minutes at room temperature. The testes were then washed 3 times in PTX (0.3% Triton-X100 in PBS). Incubated in blocking solution PBTX (5% BSA in PTX) and incubated with an appropriate dilution of primary antibodies in for overnight. Samples were washed 3 times in PTX followed by a 2-hour incubation at room temperature with Alexa dye-conjugated secondary antibodies Invitrogen) at 1:200 dilution in PBTX, and a final set of wash in PTX. The samples were mounted with a drop of Vectashield® (Vector Laboratory Inc., USA). For visualizing the nucleus, the samples were incubated with 0.001% Hoechst-33342 (Sigma Chemical Co. USA) for 20 minutes post the entire immunostaining protocol. Then the samples were washed with PTX and mounted as mentioned above. The following primary antibodies were used: rat anti-Vasa (1:50; Developmental Studies Hybridoma Bank (DSHB); developed by A. Spradling, Carnegie Institution for Science, USA), mouse anti-Armadillo (1:100; DSHB; E. Wieschaus, Princeton University, USA), rabbit anti-phospho-Histone-3 (1:4000, Santa Cruz Biotechnology); mouse anti-cyclin A (DSHB, C.F. Lehner, University of Bayreuth, Germany), rabbit cyclin-E (1:100, Santa Cruz Biotechnology).

### Quantification of germline cell death (GCD)

For detection of GCD, testes were stained with Lysotracker RedDND-99 (Life Technologies) in PBS for 30 min before paraformaldehyde fixation. For further details refer to a supplemental method, section 11, and the data is presented in Table S1.

### Determination of GSC mitotic index and the distribution of mitotically active germline cysts

We noted that staining with Lysotracker before fixation with 4% PFA led to a significant drop (by approximately 50%) in the PH3-stained pool of the germline at every stage. Therefore the PH3-staining data were independently acquired without the Lysotracker staining. The GSC mitotic index was quantified by dividing the average number of PH3-positive GSCs by the average number of GSCs in that particular genotype (Table S3). The calculations used to determine GSC cycling time have been described in the supplemental method, section 13.

### *Ex vivo* imaging of testis

The protocol for ex vivo imaging of testis was adapted from a previous study (Dubey et al., 2016). Testes from 4-day old flies were dissected and placed on a poly-lysine coated glass coverslip of a glass-bottom petri dish (P4707; Sigma). For the M-phase period estimations, the testes were immersed in Schneider’s insect medium and imaged for 2 to 4 hours. For estimation of clearance time of dying cysts, the testes were stained with Lysotracker as mentioned above and then imaged in PBS for 3 to 4 hours.

### Image acquisition, analysis, and cyst profile quantification

Images were acquired using Olympus FV1000SPD laser scanning confocal microscope using 10X, 0.3 NA and 60X, 1.35 NA or Olympus FV3000SPD laser scanning confocal microscope using 60X, 1.42 NA objectives or Zeiss 510meta laser scanning confocal microscope using the 63X, 1.4 NA objective. Live imaging was performed using FV3000SPD laser scanning confocal microscope at 10X 0.40 NA or Nikon TI-E microscope at 20X 0.75 NA Multiple optical slices were collected covering the entire apical part of the testes. The images were analyzed using ImageJ® (http://fiji.sc/Fiji). The Cell-counter^TM^ plugin was used for quantification of the immunostained cysts. The Origin (OriginLab, Northampton, MA) software and Microsoft Excel (2013) was used for all statistical analysis. For estimating the cysts stained with different cell cycle phase markers, the images were thresholded at the 95 percentile according to the intensity histogram and the cysts marked above the threshold were counted as positive.

A detailed derivation of the mathematical model is presented in supplemental methods.

Key resource table is attached listing all the the antibodies, *Drosophila* stocks and reagents used.

## Acknowledgment

We thank Benny Shilo, Talila Volk, and Eli Arama, Weismann Institute, Israel; Kenneth Irvine, Waksman Institute of Microbiology, New Jersey, USA; Dorothea Godt, Univ. Toronto, Canada; Lynn Cooley, Yale School of Medicine, Connecticut, USA; Dennis McKearin, Bloomington Stock Center, Indiana, USA; Vienna Drosophila Resource Center, Austria; and Developmental Studies Hybridoma Bank, Iowa, USA; for fly stocks and other reagents. A special thanks to Prof. Shilo for critical comments and suggestions throughout the study.

## Author’s contribution

PG: Cell cycle molecule screen, cyst distribution, pH3 distribution, and germ cell death distribution in all the genotypes and time-lapse estimation of the clearance time of dying germline cysts. PG, SC, and BV: cyst distribution in wild type and cell cycle marker frequency DM: time-lapse experiment to measure the M phase. PG, SC, and KR: data analysis. NN, PG, KR, and SG: Conceptualization of the mathematical theory. PG and KR: figures composition and manuscript writing. All the authors contributed to evolving the concept.

